# Addressing dereplication crisis: Taxonomy-free reduction of massive genome collections using embeddings of protein content

**DOI:** 10.1101/855262

**Authors:** A. Viehweger, M. Hoelzer, C. Brandt

## Abstract

Many recent microbial genome collections curate hundreds of thousands of genomes. This volume complicates many genomic analyses such as taxon assignment because the associated computational burden is substantial. However, the number of representatives of each species is highly skewed towards human pathogens and model organisms. Thus many genomes contain little additional information and could be removed. We created a frugal dereplication method that can reduce massive genome collections based on genome sequence alone, without the need for manual curation nor taxonomic information.

We recently created a genome representation for bacteria and archaea called “nanotext”. This method embeds each genome in a low-dimensional vector of numbers. Extending nanotext, our proposed algorithm called “thinspace” uses these vectors to group and dereplicate similar genomes.

We dereplicated the Genome Taxonomy Database (GTDB) from about 150 thousand genomes to less than 22 thousand. The resulting collection increases the percent of classified reads in a metagenomic dataset by a factor of 5 compared to NCBI RefSeq and performs equal to both a larger as well as a manually curated GTDB subset.

With thinspace, massive genome collections can be dereplicated on regular hardware, without affecting downstream results. It is released under a BSD-3 license (github.com/phiweger/thinspace).

## Introduction

New microbial genome collections curate hundreds of thousands of genomes.^1–3^ However, many organisms tend to be overrepresented.^4^ For example, 9.5% of genomes in the Genome Taxonony Database (GTDB)^3^ belong to *E. coli*. This redundancy complicates downstream analyses. For example, to assign reads to taxa, a k-mer index is required,^5^ and this index can only be computed on a large compute cluster for genome collections of even moderate size. One way to approach this problem is to dereplicate the collection, i.e., to remove copies of similar genomes.^6,7^ Two genomes are typically considered similar if their average nucleotide identity (ANI) is above some threshold, such as 0.95 at the species level.^8^ However, to compare each genome with all others in the collection scales quadratically with the size of the collection. We recently proposed “nanotext”, a method to represent genomes based on their protein domains, just like documents can be represented by the words they contain.^9^ Each genome is represented as an “embedding”, an n-dimensional vector of numbers. These vectors can be used as direct input to many dimensionality reduction and clustering algorithms. Here we propose “thinspace”, an algorithm that can dereplicate vast collections with millions of genomes on standard hardware without affecting task performance on read-based taxon assignment.

## Results

We dereplicated the entire GTDB with 150 thousand genomes down to 20 thousand (7.5%) on a regular laptop in under a day using a new dereplication algorithm called “thinspace”. The algorithm starts from a collection of genome sequences and precomputed embeddings. Each embedding is a latent representation of a genome’s protein content in a 100-dimensional vector of numbers.^9^ The algorithm returns the dereplicated input collection. Thinspace uses a divide-and-conquer approach and proceeds in two main steps. First, genome vectors are grouped using an unsupervised, density-based clustering method, where similar genomes – i.e., genomes with overlapping protein content – end up in the same cluster. Because genomes are represented in few dimensions, this clustering scales to potentially billions of data points.^10,11^ Second, within each cluster, pairwise nucleotide distances are computed for each genome against all others. The resulting distance matrix describes a graph, where an edge connects two vertices (genomes) if their pairwise ANI exceeds 0.95, a threshold commonly applied to discern species-level groups.^8,12^ For each connected component of this graph, the genome with the largest N50 is chosen as representative in the resulting, dereplicated collection. Large clusters are a resource constraint (e.g. a single cluster with 13,876 members of *E. coli* in the GTDB). Therefore, clusters that contain more than 500 genomes are processed in batches and iteratively recombined until a stable number of representatives is found. The entire cycle is repeated twice to increase the number of dereplicated genomes (25% reduction in the second iteration, less then 5% thereafter).

Dereplication efficiency is dependent on the quality of the clustering of similar genomes during the first step of the thinspace algorithm. The identified clusters contain homogenous taxa, and taxa rarely spread across cluster boundaries. We quantified this at the species (genus) level, with a *Rand* score of 0.65 (0.71) and a *mutual information* score of 0.76 (0.80), where 0 indicates a random allocation of point labels to clusters and 1 means perfect separation.

We validated our approach on a metagenomic read classification task for 20 recently sequenced biogas samples, comparing the collection of dereplicated genomes from thinspace (n=21,798) against a manually created, high-quality subset of genomes from the GTDB (n=24,706), hereafter called “GTDB 25k”.^4^ Both approaches perform near-identical on this task with the same percentage of long reads classified (GTDB 25k: 39.2% ± 13.6%, thinspace: 39.4% ± 13.7%, Figure 1), and they slightly underperform a much larger collection of GTDB genomes (n=54,000, 40.9% ± 14.2%, p=1, Bonferroni-corrected ANOVA).^13^ The classification based on the NCBI RefSeq collection retrieves only about one fifth of those taxonomic hits (8.5% ± 9%, p<0.001, Bonferroni-corrected ANOVA). For short reads, these results are replicable, with a larger classification variance due to less information per read (Figure 1).

**Figure 1:**
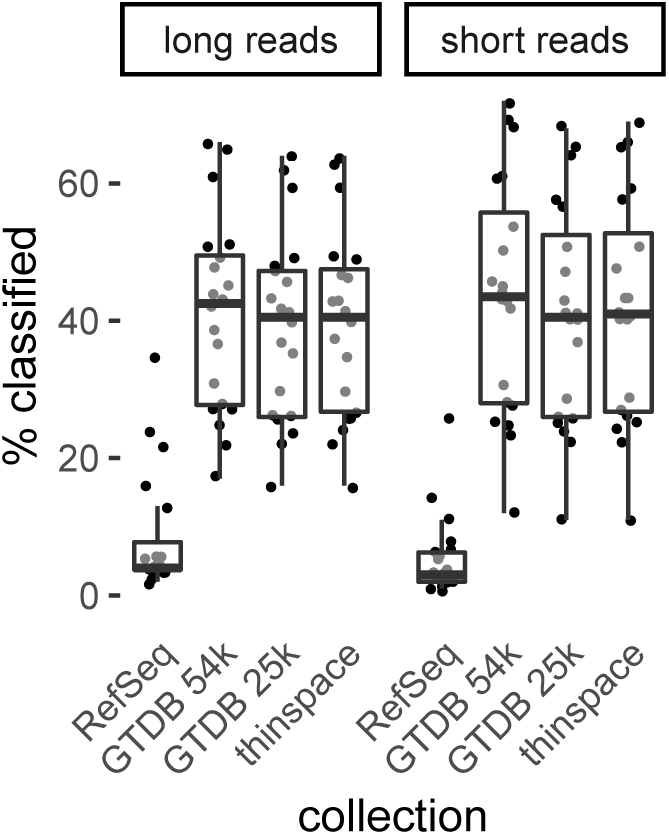
Performance of indices used in taxonomic read classification and built from different collections of genomes: NCBI RefSeq, the Genome Taxonomy Database (“GTDB 54k”), a manual dereplication of the GTDB (“GTDB 25k”)^4^ and the dereplicated collection created using thinspace. Performance was assessed on short (Illumina) and long (Nanopore) reads generated from 20 metagenomic biogas samples. As has been observed previously,^13^ all GTDB-based collections increase the number of classified reads dramatically compared to NCBI RefSeq (p<0.001). Results are similar for long and short reads. Surprisingly, when one reduces the number of reference genomes to about one representative genome per species, the percentage of reads classified does not decrease significantly, neither by manual (“GTDB 25k”^4^) nor automatic curation (thinspace).

We compared the performance of thinspace to the current standard dereplication technique of all-vs-all distance computation. Popular examples include dRep,^6^ which estimates the (nucleotide) distance between genomes with an efficient k-mer based method called Mash,^14^ as does our algorithm at the cluster level. For performance, there are two main considerations: First, the amount of memory required to store a genome (representation) limits the number of genomes that can be compared simultaneously to one another. Second, the number of pairwise comparisons determines the number of CPU hours required for the distance computations.

Thinspace uses the nanotext genome representation of a genome as a vector of numbers^9^ and requires 2 KB per genome (for a vector of 100 dimensions with floating-point numbers). Mash requires at least five times more to represent a genome, with about 10 KB for 1,000 32-bit hash integers, depending on the “sketch size”^14^ (Table 1). To compare each genome in the GTDB to all others, 10.63 billion distance calculations in 48.19 CPU hours are needed. The divide-and-conquer approach of thinspace reduces this to 0.41 billion comparisons in 1.86 CPU hours (3.85%, Table 1). The hashing of all genomes by Mash is required in both approaches.

**Table 1:** Required resources for genome dereplication of the entire Genome Taxonomy Database (GTDB, about 150 thousand genomes). Compared are an all-vs-all k-mer based approach (Mash, using *MinHash* genome sketches)^14^ and the thinspace algorithm, which relies on nanotext genome embeddings.^9^ The latter drastically reduces the required resources and can thus scale to far larger collections with millions of genomes.

| metric | unit | all-vs-all | thinspace |
| --- | --- | --- | --- |
| memory per genome | kilobyte | >10 | **2** |
| calculate distance matrix | CPU hours | 48.19 | **1.86** |
| number of comparisons | billion | 10.63 | **0.41** |

## Discussion

Microbial genome collections are growing towards a million genomes and beyond. Using these vast collections in downstream tasks is often impossible, due to the equally large compute resources required. We solve this problem by first representing genomes as vectors of numbers, and then clustering and dereplicating them in a divide-and-conquer strategy, which reduces the computational requirements dramatically. Common “batch” strategies, in which the genome collection is randomly split into subsets,^2^ are avoided. Our algorithm does not require a priori taxonomic knowledge of the input genomes. The result performs equally well compared to manual dereplication on a common classification task.

Thinspace performs two computations that the all-vs-all method does not: First, each genome needs a protein domain annotation, and all annotations are used to train the nanotext genome model.^9^ Second, the genome embeddings have to be clustered. Thinspace uses the HDBSCAN algorithm, which scales to billions of data points.^10,11^ The clustering performance can be further augmented by first reducing the dimensions of the embedding vectors, for which thinspace provides an option using the UMAP algorithm.^15^ This upfront cost has advantages compared to an approach purely based on nucleotide distance: Nanotext vectors can correctly represent even incomplete genomes,^9^ which Mash cannot.^6^ Also unlike Mash, nanotext can reliably calculate pairwise genome distances below an ANI of 0.8.^9,14^ In conclusion, thinspace can dereplicate millions of even incomplete genomes over large distances on standard hardware in a reasonable time.

## Methods

For genome embeddings, we used the nanotext library (v0.1, github.com/phiweger/nanotext) with the “core” model which emphasises core genes for genome comparison. The resulting vectors of numbers were clustered using HDBSCAN.^15^ To assess the quality of the clustering, we use the *Rand* score (adjusted)^16^ and the *mutual information* score (adjusted)^17^ as implemented in the sklearn library (v0.21.3, scikit-learn.org). Clusters were dereplicated using custom scripts developed by Ryan Wick (github.com/rrwick/Assembly-Dereplicator) as was the reformatting of the results (github.com/rrwick/Metagenomics-Index-Correction) to be compatible with index generation by Centrifuge.^5^

We compared precomputed indices from NCBI RefSeq and the GTDB^13^ to those we generated for dereplicated GTDB subsets, one that was manually curated^4^ and one generated by thinspace. After assigning reads of at least 150 bases to taxa, the Centrifuge output was filtered to above a quality score of 250. The biogas metagenomes are available from the European Nucleotide Archive (ENA) under the project accession PRJEB34573.

## Contributions

AV designed and implemented the thinspace algorithm. AV, MH and CB designed the benchmark experiments. CB performed metagenomic sequencing. AV wrote the draft of this article, on which MH and CB provided extensive advice.

